# Multiplexed direct RNA sequencing using RB-dRNAseq with RNA barcodes

**DOI:** 10.64898/2026.05.14.725048

**Authors:** Zhuo-Xing Shi, Qing-Pei Huang, Ying-Feng Zheng, Feng Zhang, Xiao-Ying Fan

## Abstract

Nanopore direct RNA sequencing (dRNA-Seq) simultaneously resolves RNA sequences, poly(A) tail lengths, and base modifications at the single-molecule level. Here, we developed RB-dRNAseq, a multiplexed dRNA-Seq method using RNA-barcoded reverse transcription adapters (RTAs) compatible with Oxford Nanopore Technologies (ONT) workflows, enabling highly efficient sample demultiplexing based solely on basecalled sequences. On the RNA004 platform, RB-dRNAseq achieved >99.5% demultiplexing accuracy, with 81% of reads assigned in 6-plex samples. This represents a 16%-24% relative increase in read recovery compared to conventional DNA barcoding methods that rely on basecalling-based demultiplexing. To evaluate the sensitivity of RB-dRNAseq, we performed sequencing with low-input samples, demonstrating that even with 1 ng of total RNA, gene expression (r=0.84) and m6A modification (r=0.91) profiles remained highly consistent with high-input benchmarks. We further explored the feasibility of RB-dRNAseq in ultra-low input scenarios by applying it to mouse zygotes and 2-cell stage embryos. The method successfully captured dynamic shifts in gene isoform expression and multiple RNA modifications (m6A, m5C, pseU, and Inosine) during zygotic genome activation (ZGA), uncovering site-specific regulatory patterns.

## Introduction

The nanopore sequencing platform enables direct sequencing of native RNA(Stark et al. 2019; Jain et al. 2022; Monzó et al. 2025), concurrently resolving base sequence(Xin et al. 2021; Zhang et al. 2021), poly(A) tail length(Soneson et al. 2019; Parker et al. 2020; Roach et al. 2020), and RNA base modifications for individual RNA molecules(Garalde et al. 2018; Begik et al. 2021; Lucas and Novoa 2023; Lucas et al. 2023; Liu and Conesa 2025). However, the nanopore dRNA-Seq methods for low-input samples remain scarce, primarily due to several technical challenges(Stark et al. 2019; Pardo-Palacios et al. 2024; Liu and Conesa 2025). First, RNA extraction protocols for trace samples are underdeveloped. Second, current library preparation and sequencing workflows demand high RNA input(Pardo-Palacios et al. 2024). For instance, the recommended input for dRNA-Seq using the ONT RNA004 flow cell is 1 μg total RNA or 300 ng poly(A)+ RNA, far exceeding the RNA content of most low-input samples and limiting its application.

Multiplexing samples using barcoded Reverse Transcription Adapters (RTAs) offers a viable strategy to increase the effective starting material and thus ensure sufficient sequencing yield in dRNA□Seq(Foord et al. 2023). However, ONT has not yet officially released a barcoding method for dRNA-Seq. A key challenge is that the RTA sequence is DNA, and DNA barcodes are co-sequenced with the RNA strand, making them difficult for RNA basecalling software to recognize, thus hindering their use in dRNA-Seq. Recent academic developments include several multiplexed dRNA-Seq techniques using DNA-barcoded RTAs. For example, Smith et al. (2020) proposed DeePlexiCon, which designs a 20 bp DNA barcode within the RTA and utilizes deep learning models trained on the current signal of the barcode sequences for demultiplexing. On the RNA002 platform, DeePlexiCon classified 60% of reads with 99.9% accuracy using 4 barcodes(Smith et al. 2020). Subsequently, Toorn et al. released WarpDemuX, employing a similar DNA-barcoded RTA scheme and extending the current signal-based deep learning demultiplexing model to 12 barcodes, further increasing the sample throughput(van der Toorn et al. 2025). Additionally, SeqTagger improved upon DeePlexiCon by expanding barcode diversity and adapting the approach to the RNA004 platform, and it performs demultiplexing via basecalling with a custom-trained Bonito model(Pryszcz et al. 2025). While DeePlexiCon, WarpDemuX, and SeqTagger provide feasible barcoded dRNA-Seq solutions and allow researchers to use pre-trained models, their flexibility is limited. Customization requires additional deep learning model training, increasing the barrier to use. Furthermore, these methods necessitate retraining following major ONT upgrades to reagents, flow cells, or basecalling algorithms.

Here, we developed RB-dRNAseq, which employs RNA-barcoded RTAs compatible with the ONT dRNA-Seq library prep, enabling high-precision demultiplexing post-sequencing based solely on the basecalled sequence. We demonstrated that the RNA barcoding strategy significantly increased the demultiplexing efficiency compared to DNA barcoding, and the performance improved from RNA002 to RNA004 platform. We show that RB-dRNAseq accurately capture the isoform expression, poly(A) tail length and epitranscriptome information across different total RNA input amounts. Finally, we applied RB-dRNAseq to mouse zygote and 2-cell stage embryos, resolving isoform expression and modification dynamics at the single-molecule level during zygotic genome activation (ZGA).

## Results

### High accuracy and specificity of demultiplexing DRS data using the RNA barcoding strategy

For RNA-barcoded RTA design in RB-dRNAseq, we developed OligoA (5’-10DNA-24RNA-3DNA-3’) where the 20 RNA barcode bases are flanked by DNA sequences, and OligoB is entirely DNA (Fig. 1A). OligoA and OligoB are synthesized separately and annealed to form the complete barcoded RTA (Supplementary Table 1). This single-stranded RNA barcode design reduces synthesis costs and the three DNA bases at the 3’ end of OligoA ensure compatibility with the ONT ligation step and mitigate RNA degradation risk. We replaced the standard RTA component with our RB-barcoded RTA, all other library preparation and sequencing steps strictly followed the ONT protocol when using total RNA input (Fig. 1B).

**Figure 1.**
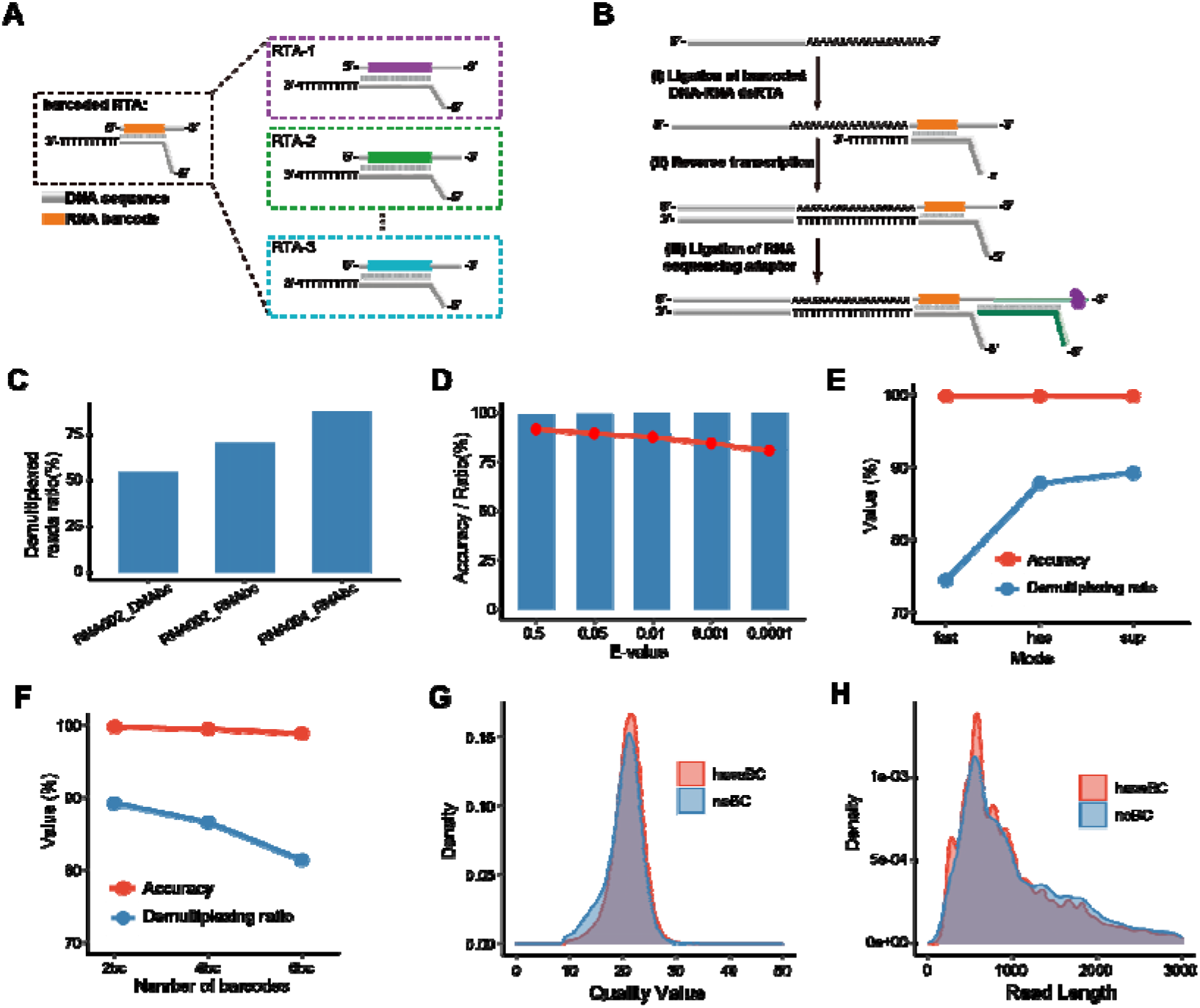
Schematic and performance evaluation of RB-dRNAseq. **(A)** Schematic of barcoded RTA design. **(B)** RB-dRNAseq library preparation workflow. **(C)** Bar plot comparing demultiplexing rates of RNA002 DNA barcode, RNA002 RNA barcode, and RNA004 RNA barcode strategies (hac basecalling mode, e-value<0.05). **(D)** Line plot (red, demultiplexing rate) and bar plot (blue, accuracy) for RNA004 RNA barcode under different e-value thresholds (super basecalling mode). **(E)** Line plots comparing demultiplexing rate and accuracy for RNA004 RNA barcode under different Dorado basecalling modes (e-value<0.05). **(F)** Line plots comparing demultiplexing rate and accuracy for RNA004 RNA barcode with different numbers of barcodes (sup basecalling mode, e-value<0.05). **(G)** Read length distributions for reads with demultiplexed barcodes vs. undemultiplexed reads (sup basecalling mode, e-value<0.05). **(H)** Sequencing quality (Q-score) distributions for reads with demultiplexed barcodes vs. undemultiplexed reads (sup basecalling mode, e-value<0.05).

We first tested the RNA barcode strategy on RNA002 and this largely improved the reads demultiplexing ratio (70.44%) compared with the DNA barcodes (54.51%, Fig. 1C). The ratio of demultiplexed reads further improved on RNA004 (87.77%) using the RNA barcode strategy (Fig. 1C, Supplementary Table 2). We examined the impact of barcode alignment e-value thresholds on demultiplexing rate and accuracy. As the e-value threshold became stricter, the accuracy changed minimally (99.14%-99.88%), but the demultiplexing rate gradually decreased (Fig. 1D, Supplementary Fig. 1A-B, Supplementary Table 2). We finally selected an e-value threshold of 0.05 for downstream analysis.

We further evaluated the impact of different ONT Dorado basecalling modes on demultiplexing rate and accuracy. The Dorado sup mode significantly increased the barcode demultiplexing rate compared to the fast mode, which the accuracy slightly decreased (Fig. 1E, Supplementary Fig. 1C-D, Supplementary Table 2). Additionally, increasing the number of barcode types from 2 to 6 also caused the demultiplexing rate decreased from 89.2% to 81.4% (Fig. 1F, Supplementary Fig. 1E-F, Supplementary Table 2). We checked whether the failed reads (without barcode sequence recognition) were caused by the sequencing quality or the read length, but neither of the factors exhibited differences between barcoded reads and non-barcoded reads (Fig. 1G-H). This might be caused by defects of the current basecalling algorithm at the barcode region. Future optimization of the basecalling algorithm may improve the overall demultiplexing rate, supporting the use of more barcode types.

### Validation of RB-dRNAseq across different total RNA input

To evaluate the performance of RB-dRNAseq across varying RNA input amounts from different species, we extracted total RNA from human HEK293T cells and mouse embryonic stem cells (mESCs), respectively. We used 1 ng, 10 ng, 100 ng, 500 ng of total RNA as the starting material, corresponding to four different barcodes (Fig. 2A). Barcode demultiplexing demonstrated that RB-dRNAseq accurately recapitulated the output read counts corresponding to different input amounts (Fig. 2B, Supplementary Table 3). Quality control analysis showed stable read length distributions across input levels (Fig. 2C), and there were no significant difference between the average poly(A) tail lengths detected in the low-input and high-input RNA samples (Fig. 2D).

**Figure 2.**
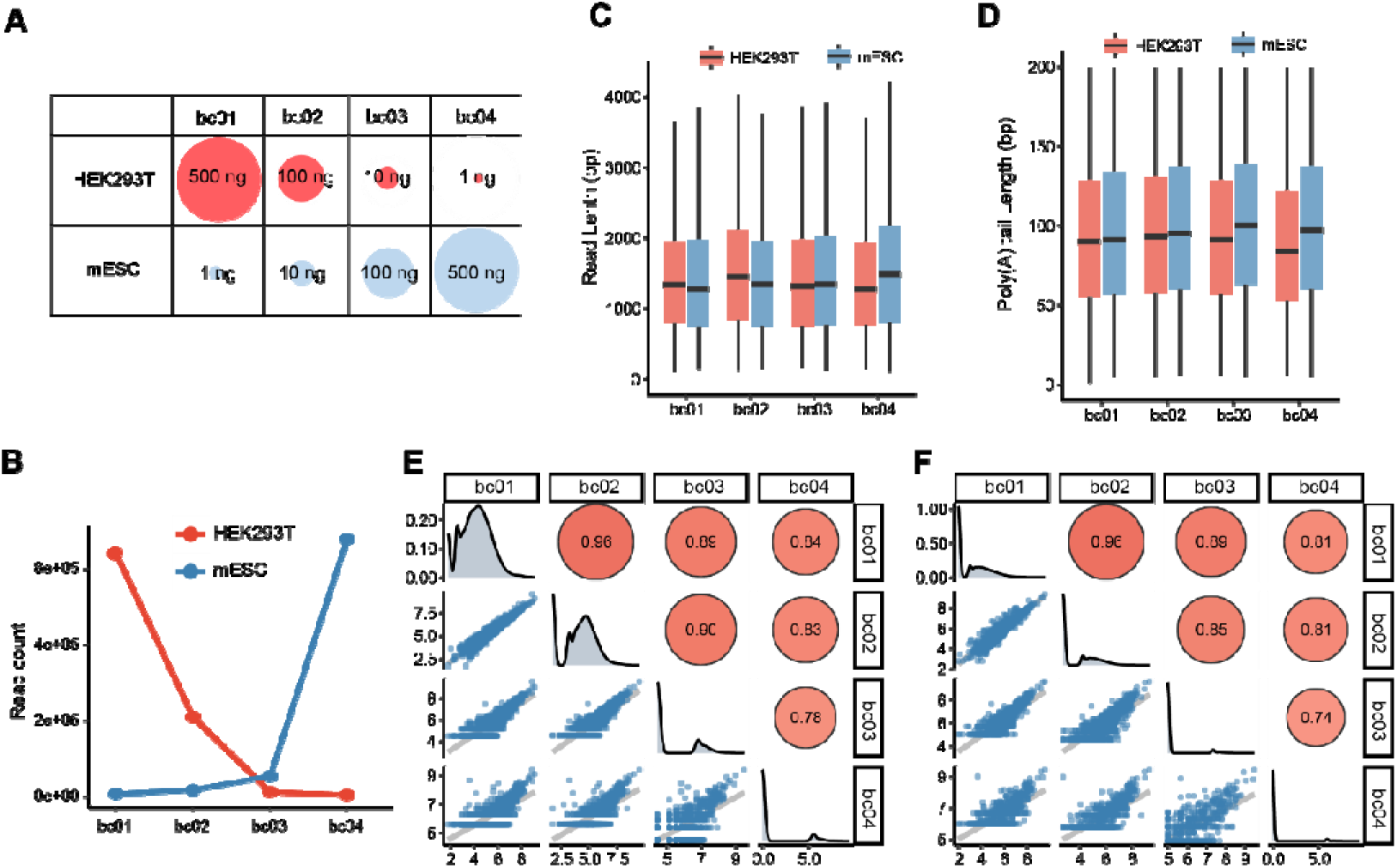
Performance of RB-dRNAseq with different total RNA input. **(A)** Experimental design for multiplexed RB-dRNAseq with varying input amounts. **(B)** Line plot comparing sequencing read counts across input amounts. **(C)** Boxplot of read length distributions across input amounts. **(D)** Boxplot of poly(A) tail length distributions across input amounts. (E-F) Scatter plot showing gene expression **(E)** and isoform expression **(F)** correlation between different HEK293T total RNA input indicated by the RTA barcodes.

Then we compared the performance in gene and transcript detection by RB-dRNAseq when starting with different amounts of input RNA. Both the number of genes and isoforms largely increased from 1ng to 500ng total RNA as input (Supplementary Fig. 2A-B). This should attribute to lower sample input leading to fewer sequencing output reads. We confirmed that the number of detected genes largely increased with the number of obtained reads under current shallow sequencing depth (Supplementary Fig. 2C-D). For 1ng total RNA, we detected 1828 genes, 1371 transcripts in human and 2531 genes, 1791 transcripts in mouse (Supplementary Table 3). Although the number of genes and isoforms detected by 1ng total RNA is only one-tenth of that detected by 500ng total RNA, the correlation between the gene expressions obtained from the two groups is still high (correlation coefficient value 0.81-0.86, Fig. 2E-F, Supplementary Fig. 3A-B). This indicates reliable transcript quantification in low-input samples using RB-dRNAseq.

As RB-dRNAseq also reports RNA modifications at single-base resolution, we evaluated the accuracy of m6A detection in each group. Similar to expression detection, the number of detected m6A sites highly relies on the amount of input sample (Supplementary Fig. 4A-B). Metagene distribution profiles of m6A sites all showed enrichment in the 3’ UTR, consistent to the classical m6A distribution (Fig. 3A-B). However, the 100 and 500ng groups displayed higher density of m6A sites in the 3’UTR than the 10 and 1ng groups, indicating that the low-input samples would prefer lose the capture of m6A in this region. Interestingly, the quantification of the m6A modification level is not significantly affected by the initial amount of the input RNA, as the correlation values among different groups of samples did not changed obviously (Fig. 3C-D). These data robustly demonstrates accurate capture of m6A sites and precise quantification of the modification levels in low-input samples using RB-dRNAseq.

**Figure 3.**
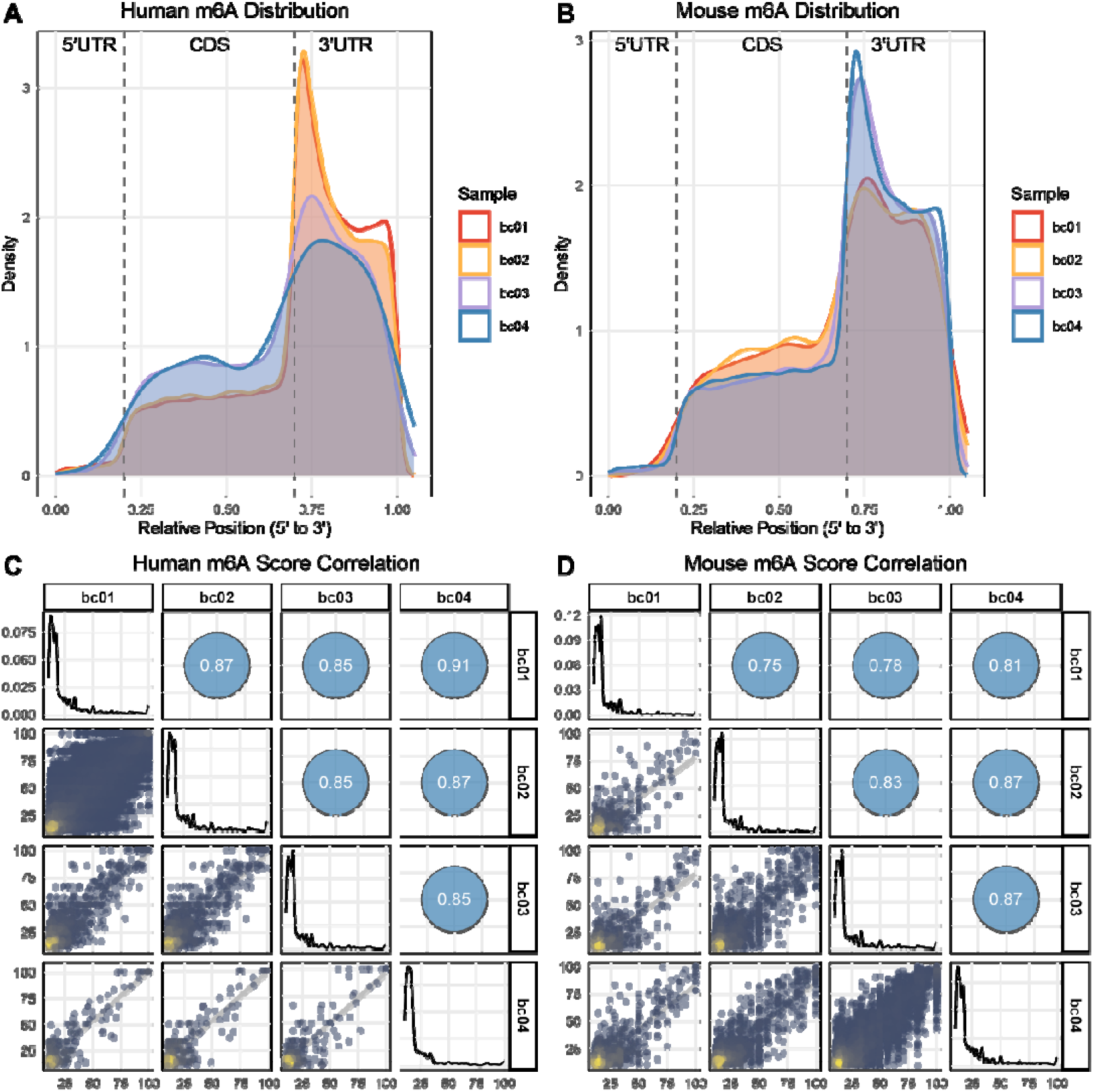
Comparison of m6A detection by RB-dRNAseq across different total RNA input. (A-B) Metagene distribution of m6A sites for HEK293T **(A)** and mESC **(B)** across different total RNA input indicated by the RTA barcodes. (C-D) Scatter plot showing the correlation of modification level of each m6A site between different HEK293T **(C)** and mESC **(D)** total RNA input indicated by the RTA barcodes.

### RB-dRNAseq in low-input cell samples

dRNA-Seq has been only applied for large amount of total RNA or poly(A) RNA in previous studies. We proved that RB-dRNAseq accurately capture expression and modification of as few as 1ng total RNA. However, in practical applications, such amount of samples are generally not subjected to RNA extraction. So added cell lysis and RNA capture step before ligating RTAs and applied the workflow in mouse zygote and 2-cell embryos. We started with as few as 1 embryos in each sample replicate. Meanwhile, we added spike-in RNA in the library to maintain the activity of the sequencing holes and ensure sufficient sequencing yield (Fig. 4A, Supplementary Table 4).

**Figure 4.**
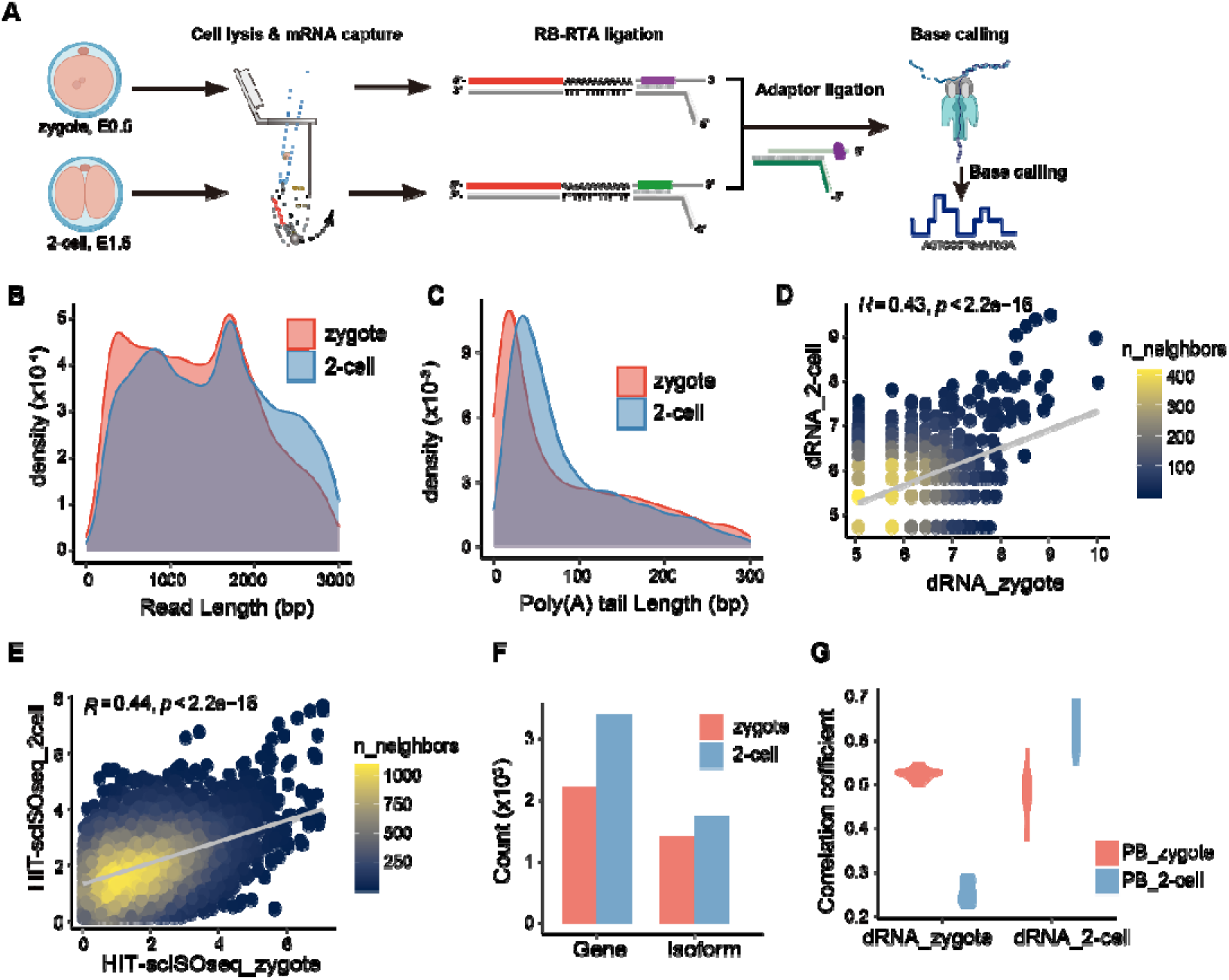
RB-dRNAseq results for mouse zygote and 2-cell samples. **(A)** Schematic of RB-dRNAseq for mouse zygote and 2-cell embryos. (B-C) Distribution of read lengths **(B)** and poly(A) tail lengths **(C)** for zygote and 2-cell embryos. **(D)** Bar plot showing number of genes and isoforms detected by RB-dRNAseq in zygote and 2-cell embryos. (E-F) Scatter plot of gene expression correlation between zygote and 2-cell samples using RB-dRNAseq **(E)** and HIT-scISO-seq **(F). (G)** Violin plot showing distribution of gene expression correlation values between RB-dRNAseq and HIT-scISOseq at the single-cell level for zygote and 2-cell samples.

Due to too few reads generated from several batches of samples, we merged reads from the same stages and compared the transcript lengths and poly(A) tail lengths at the zygote and 2-cell stages. The 2-cell embryos contains longer RNAs with longer poly(A) tails (Fig. 4B-C), which is consistent to previous studies (e.g., Liu et al.(Liu et al. 2023)). Further mapping revealed 2218 genes and 1405 RNA isoforms in zygotes, and 3401 genes and 1759 isoforms in 2-cell embryos (Fig. 4D). Integration with our previously published HIT-scISOseq data showed higher concordance in gene expression between RB-dRNAseq and HIT-scISOseq for the same stages of samples (Fig. 4E-G), validating the reliability of RB-dRNAseq data for expression quantification of low-input cell sample.

### Isoform expression and RNA modification changes during ZGA

The fact that ZGA serves as the foundation for the emergence of life has received widespread attention. Previous single-cell methods enable us to investigate gene expression changes during this process. Due to limitations in the current single-cell epitranscriptome detection methods, little is known about how RNA modification changes during ZGA. With RB-dRNAseq from zygote and 2-cell embryos, we focused on differences in multiple RNA modifications at RNA isoform resolution. We performed joint detection of multiple RNA modifications (m6A, m5C, pseU, Inosine) on single molecules using ONT’s Dorado basecaller (see Methods). Subsequently, the ONT modkit tool was used to map read-level methylation information to RNA isoforms, yielding single-base resolution modification information for each gene isoform. For all four kinds of modifications analyzed, the modified sites in zygotes showed higher modification levels than those in 2-cell embryos (Fig. 5A), consistent with previous reports (e.g., Wu et al.(Wu et al. 2022)). Further comparison of differentially modified sites (DMS, difference exceeds 5%) returned more sites exhibiting reduced modification levels in 2-cell embryos. Specifically, a total of 2796 DMS were identified, and the number of DMS for m6A and pseU was significantly higher than for m5C and Inosine (Fig. 5B). Interestingly, we observed different modification changes for different sites on the same transcript. Take the *ENSMUST00000114431* from *Btg4* as example, m6A and Inosine levels were generally low on this transcript in both group of samples; the zygote showed higher pseU level at pos167 than the 2-cell embryo, while the opposite trend was observed at pos437 and pos520; the 2-cell embryo exhibited increased m5C level at pos517 and decreased at pos648 (Fig. 5C). Importantly, we can determine how same and different modifications are coordinated on the same RNA molecule (Fig. 5D). These results fully demonstrate RB-dRNAseq’s capability to resolve heterogeneity of multiple RNA modifications across samples at isoform and single-base resolution.

**Figure 5.**
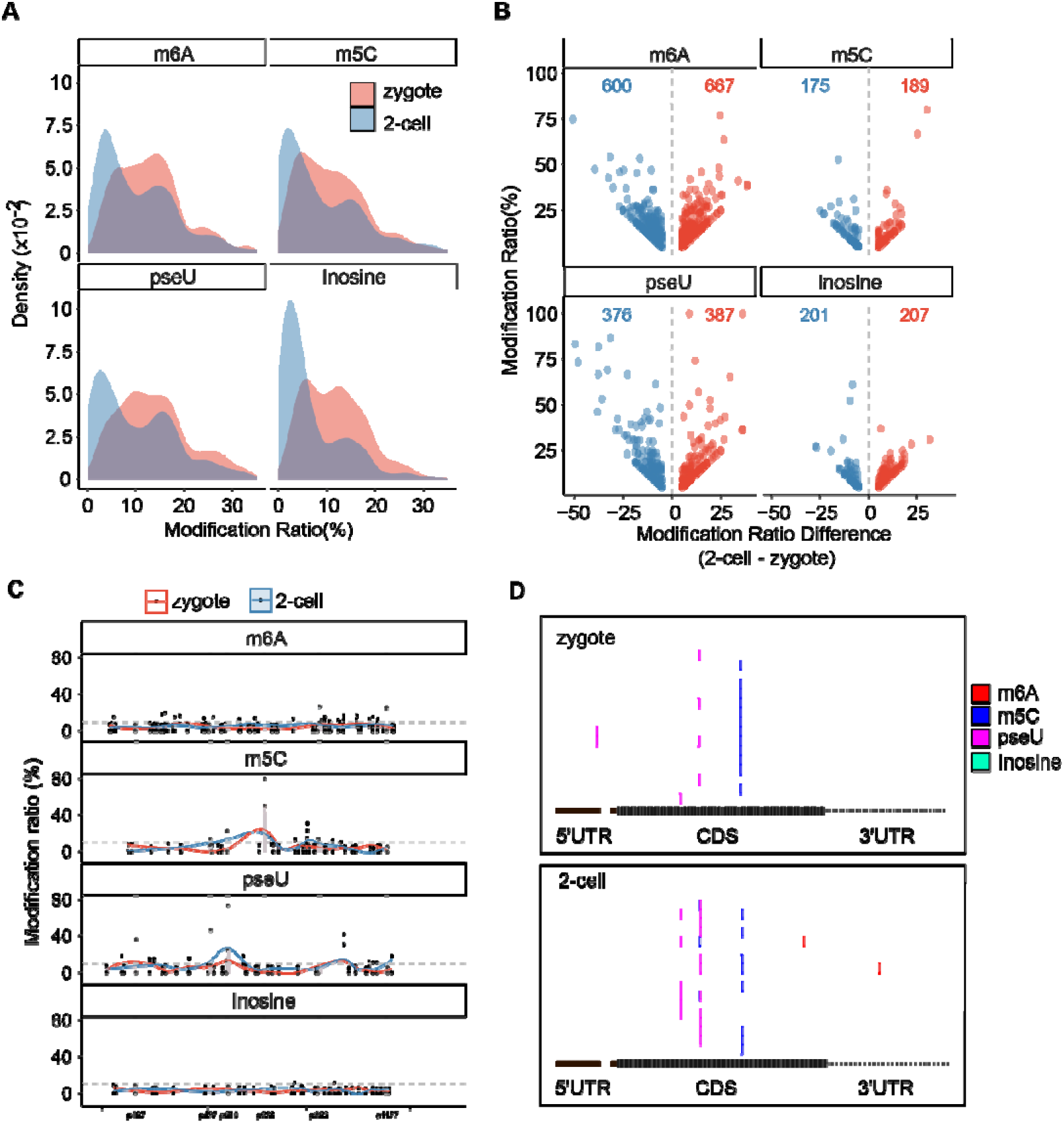
RNA modification analysis at single molecular and single base resolution. **(A)** Density distribution of modification levels of detected sites in the isoforms. **(B)** Volcano plots showing DMS (2-cell vs zygote) for different modification types. **(C)** Methylation abundance profiles for different modification types across the *Btg4* transcript (*ENSMUST00000114431*) in zygote and 2-cell samples. **(D)** Single-molecule read visualization of co-occurrence of different modification types at specific sites on the *Btg4* transcript (*ENSMUST00000114431*) for zygote and 2-cell samples.

## Discussion

Multiplex sampling has been the major challenge for dRNA-Seq. We developed RB-dRNAseq, which addresses the core limitation of DNA barcodes in dRNA-Seq - unrecognizability by basecalling software, by designing the barcode sequence as RNA. Meanwhile, we designed the DNA-RNA chimera sequence to reduce costs while ensuring compatibility with ONT workflows via the 3’ DNA anchor. Compared to DNA barcoding methods relying on current signal deep learning (e.g., DeePlexiCon), RB-dRNAseq enables demultiplexing through simple sequence alignment, eliminating the complexity of model training and platform adaptation, while significantly improving accuracy (>99.5%) and multiplexing capacity (>81% demultiplexing rate at 6-plex). RB-dRNAseq maintains stable transcript quantification (gene expression correlation r=0.84) and modification detection (m6A abundance correlation r=0.91) even at the ultra-low input of 1 ng, with no significant bias in read length or poly(A) tail distribution. Combined with a direct post-lysis library construction strategy for single cells/embryos, the method was successfully applied to study mouse early embryonic development. The detected transcript expression patterns during ZGA were highly consistent with HIT-scISOseq data, validating data reliability. In this way, we are able to integrate multidimensional information, including isoform expression, poly(A) tail length, and multiple modifications concurrently on single molecules, revealing coordinated changes during the ZGA process (e.g., on *Btg4*).

However, there are also some limitations for the current version of RB-dRNAseq. First, the 24 nt single-stranded RNA barcodes used in RB-dRNAseq have relatively high synthesis costs. Given the current 6-plex RNA barcode edit distances of 12-18 (Supplementary Table 5), future optimization could reduce barcode length, thereby lowering costs. Additionally, the demultiplexing ratio could be improved, especially for more sample numbers. The DNA sequences flanking the RNA barcode may cause basecalling errors. Future optimization of algorithms or targeted model training is needed to improve demultiplexing rates. Second, the current sequencing yields for low-input samples (including both total RNA and cells) are quite low. We tried to add high amounts of spike-in RNA to ensure the activity of the sequencing holes, thus to generate more total reads. But the proportion of reads from target samples is relatively low. The best way to solve this problem is to enhance the efficiency of ONT sequencing. This may extend the application of RB-dRNAseq in single-cell and spatial transcriptome analysis, and rapid pathogen detection, etc.

## Methods

### Animals and single blastomere collection

We used 8 to 12 week-old C57BL/6J female mice and DBA/2NCrl male mice in the experiment. The female mice were first injected with 7.5 IU of pregnant mare’s serum gonadotropin (PMSG) (Ningbo SanSheng Biological Technology, Cat. 110044564), and with 7.5 IU of human chorionic gonadotropin (hCG) (Ningbo SanSheng Biological Technology, Cat. 50030248) after 48 hours. After mating, the embryos of each stage were collected at defined time periods after hCG injection: 22 to 24 hours (zygote), 46 to 48 hours (late 2cell). All animal experiments were performed according to the guidelines of the Guangzhou national laboratory (Guangzhou, China). Collection of single blastomeres at each stage was carried out as previously described(Koo et al. 2020).

### mRNA capture for total RNA and cell lysate

Total RNA was extracted from HEK293T and mESC (derived from C57BL/6J mice) by RNA extraction kit (Yeasen, Cat. 19221ES)respectively. For early mouse embryos, including zygotes and 2 cells, single embryo was taken and placed into the cell lysis buffer (0.8% Triton X-100 (Sigma-Aldrich, T9284), 2 mM DTT (Sigma-Aldrich, 43815), 2U/µl of RNase inhibitor (TaKaRa, 2313A)). Vortex vigorously for 10 seconds to ensure complete lysis.

Resuspend the mRNA capture magnetic beads (Vazyme, Cat. N401), take 10 μL of magnetic beads into a 200 μL volume centrifuge tube, place it on a magnetic stand for 2 minutes, and discard the supernatant. Add 40 μL of 1× HB buffer (3× Saline-Sodium Citrate Buffer (Sigma-Aldrich, S6639), 0.05% Tween-20 (Sigma-Aldrich, 655204), 2U/µl of RNase inhibitor) to wash the magnetic beads and repeat the washing twice. Add 40 μL of 2× HB buffer (6×SSC, 0.05% Tween-20, 2U/µl of RNase inhibitor) to resuspend the magnetic beads.

Add an equal volume of the magnetic beads in 2× HB buffer to total RNA or cell lysate, mix gently, and then instantly centrifuge at low speed and place it in PCR instrument, incubate at 65°C for 5 minutes, then place it on a rotating rack at room temperature for 15 minutes. Leave it on a magnetic stand for 2 minutes and discard the supernatant. Add 40 μL of pre-cooled 1× HB buffer to resuspend the magnetic beads, centrifuge at low speed instantaneously, and then place it on a magnetic stand for 2 minutes, discard the supernatant. Repeat washing twice. Add 40 μL of pre-cooled 1× phosphate-buffered saline (PBS, Invitrogen, AM9625) to wash the magnetic beads once. Remove the supernatant thoroughly. Add 10 μL of RNase-free water, flick and resuspend thoroughly, instantly centrifuge at low speed, place it in a PCR instrument, incubate for 2 minutes at 80°C, quickly incubate on ice for 2 minutes, then place it on a magnetic stand for 5 minutes, transfer the supernatant to a new 200 μL volume centrifuge tube.

### Design and assembly of custom reverse transcription adapters (RTAs) containing Barcode

RTA is made of the forward chain RTA-F and the reverse chain RTA-R. The forward strand RTA-F is a DNA-RNA chimeric strand composed of DNA bases and RNA bases. Its structure is as follows: 5’-TCCN(24)TAGTAGGTTC-3’, where N(24) is a 24nt-length RNA sequence, which is the barcode region of RTA. The other bases are DNA bases. RTA-R consists of DNA bases and has a structure as follows: 5’-GAGGCGAGCGGTCAATTTTM(24)GGATTTTTTTTTTT-3’, where M(24)GGA is the reverse complementary sequence of TCCN(24) in RTA-F. The Barcode sequences are shown in supplemental table1. The RTA-F and RTA-R oligo are synthesized by Sangon Biotech (Shanghai).

The RTA-F and RTA-R of each RTA were diluted to 10 μM and 9 μM with annealing buffer (10 mM Tris-HCl, pH 7.5, 50 mM NaCl, 0.5 mM EDTA), respectively. Take 5 μL each and mix it and then denature and anneale. The reason for using excessive forward strands in RTA assembly is that excessive forward strands can ensure complete consumption of reverse strands, avoiding the reduction of RNA-RTA products after annealing, resulting in competition with RTA during subsequent reverse transcription. The assembly of custom RTA was carried out according to the following PCR procedure: denaturation at 98°C for 3 minutes, decreased by 1°C per minute until 25□, and maintains at 4□. RTA with different RNA barcodes could be used for direct RNA sequencing library construction.

### Sample multiplexing and library construction

According to the ONT Direct RNA Sequencing Product Manual (SQK RNA004), the captured mRNA of each sample is ligated with RTA containing certain barcode, and undergoes reverse transcription reaction to form an RNA-DNA hybrid strand. The RNA/DNA hybrid is then ligated with RNA ligation adapter (RLA) to form a complete sequencing structure.

Brief steps are as follows: For each sample, an RTA containing a specific barcode is ligated with its RNA using NEBNext rapid ligase (NEB M0202M) at room temperature for 30 minutes. Reverse transcription was then performed using SuperScriptIV reverse transcriptase (ThermoFisher, 18090050). The prouducts of reverse transcription with different RTAs were mixed together and then purified using 2× RNA Clean XP beads (Beckman, A63987). The purified product was ligated with the RLA linker using NEB Nextflash ligase (NEB M0202M) at room temperature for 30 minutes. The reaction mixture was purified using 0.6× RNA Clean XP beads (Beckman, A63987), washed twice with buffer (WSB), and the product was eluted with elution buffer (EB). Then prepare the priming mixture according to the instructions (SQK RNA004) and adjust it according to the volume of the library adapted to the PromethION sequencer. Load the mixed solution on the machine into the ONT sequencing cell that passes the quality check, set the sequencing parameters in the MinKNOW software, and the sequencing lasts for more than 12 hours to obtain sufficient sequencing data.

### Basecalling and demultiplexing

After obtaining barcode-labeled DRS data, use ONT’s official Dorado software and set the “--no-trim” parameter to retain the sequencing adapter, and perform basecalling on the original pod5 data. Three precision modes could be chosen: fast (fast), high precision (hac), and super-high precision (sup). Next, the blastn program was used to align the RTA barcode sequences to the DRS reads after basecalling. The alignment results are filtered based on the alignment length greater than 9, the alignment position (s.end) at the 5’ terminal of barcode greater than 5, and the e-value of the alignment. Finally, for each DRS read, the barcode with the smallest e-value in alignment was selected as the barcode of the read.

## Supporting information

Source data for Supplementary Table 2

## Data availability

All sequencing data will be publicly released upon acceptance of the manuscript for publication. Source data for Supplementary Table 2 are provided as a supplementary data file.

## Acknowledgements

X.-Y.F. was supported by grants from the Guangdong Provincial Pearl River Talents Program (2021QN02Y747), Guangzhou science and technology elite “pilot” project (SL2024A04J01788) and the Major Project of Guangzhou National Laboratory (GZNL2023A02003, GZNL2024A03001). Z.-X.S. was supported by grants from the National Natural Science Foundation of China (42107148).

## Author Contributions Statement

X.-Y.F., and F.Z. conceived and designed the project; Z.-X.S., X.-Y.F., and Q.-P.H. developed the experimental technology; Q.-P.H performed sequencing experiments. Z.-X.S., and Q.-P.H. performed the informatics analysis; Z.-X.S., Q.-P.H., and Y.-F.Z. coordinated data release and assisted with executing the pipeline. Z.-X.S., Q.-P.H., and X.-Y.F. wrote the manuscript and created the figures; Y.-F.Z., and F.Z. advised the study and revised the manuscript. All authors have read and approved the final version of this manuscript.

