## Supplementary Table1-5 and Supplementary Figure1-4 for "Multiplexed direct RNA sequencing using RB-dRNAseq with RNA barcodes"

### Supplementary Information

#### TABLE OF CONTENTS

##### Supplementary Figures


Supplementary Figure 1. Performance evaluation of RB-dRNAseq.

Supplementary Figure 2. Gene and isoform detection between different total RNA inputs.

Supplementary Figure 3. Gene expression and isoform expression correlation between different total RNA inputs.

Supplementary Figure 4. The m6A site detection between different total RNA inputs.


##### Supplementary Tables


Supplementary Table 1. The sequences of RTA oligo and barcodes.

Supplementary Table 2. The results of barcode demultiplexing rate and demultiplexing accuracy.

Supplementary Table 3. The results of barcode demultiplexing and feature detection between different total RNA inputs.

Supplementary Table 4. The results of barcode demultiplexing of low input embryos samples.

Supplementary Table 5. The edit distance matrix between RNA barcodes.

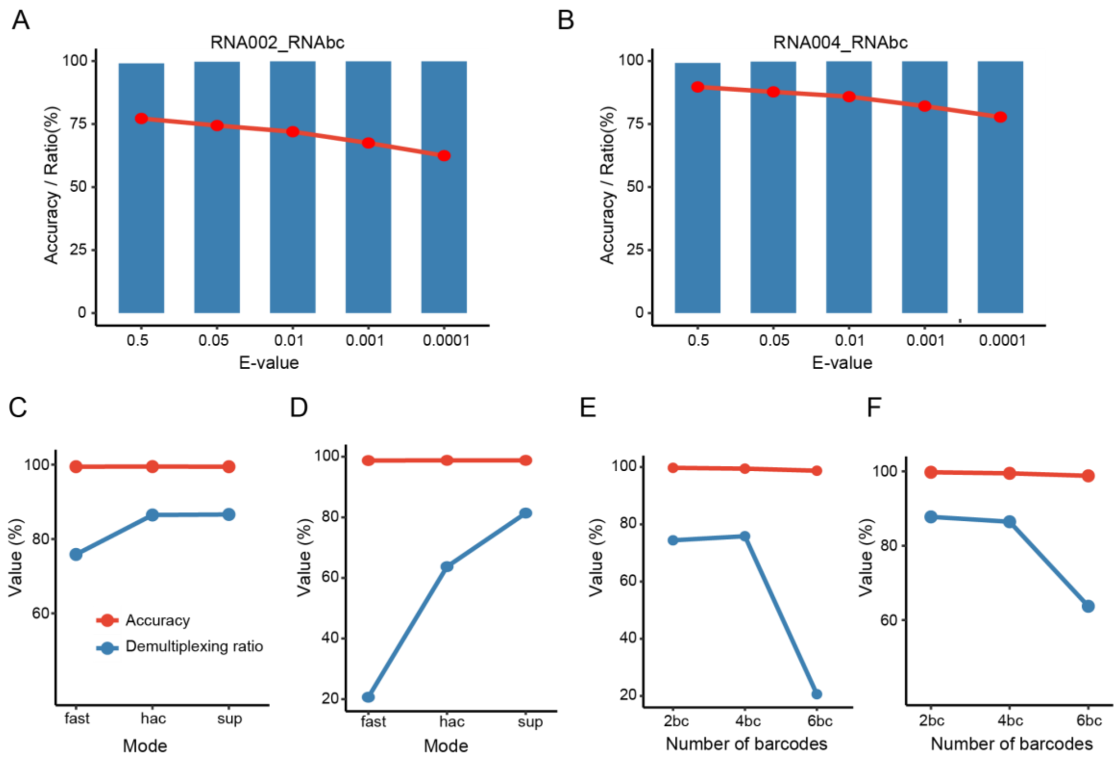


**Supplementary Figure 1. Performance evaluation of RB-dRNAseq. (A)** Line plot (red, demultiplexing rate) and bar plot (blue, accuracy) for RNA002 2 RNA barcode under different e-value thresholds (fast basecalling mode). **(B)** Line plot (red, demultiplexing rate) and bar plot (blue, accuracy) for RNA004 2 RNA barcode under different e-value thresholds (hac basecalling mode). **(C)** Line plots comparing demultiplexing rate and accuracy for RNA004 4 RNA barcode under different Dorado basecalling modes (e-value<0.05). **(D)** Line plots comparing demultiplexing rate and accuracy for RNA004 6 RNA barcode under different Dorado basecalling modes (e-value<0.05). **(E)** Line plots comparing demultiplexing rate and accuracy for RNA004 RNA barcode with different numbers of barcodes (fast basecalling mode, e-value<0.05). **(F)** Line plots comparing demultiplexing rate and accuracy for RNA004 RNA barcode with different numbers of barcodes (hac basecalling mode, e-value<0.05).

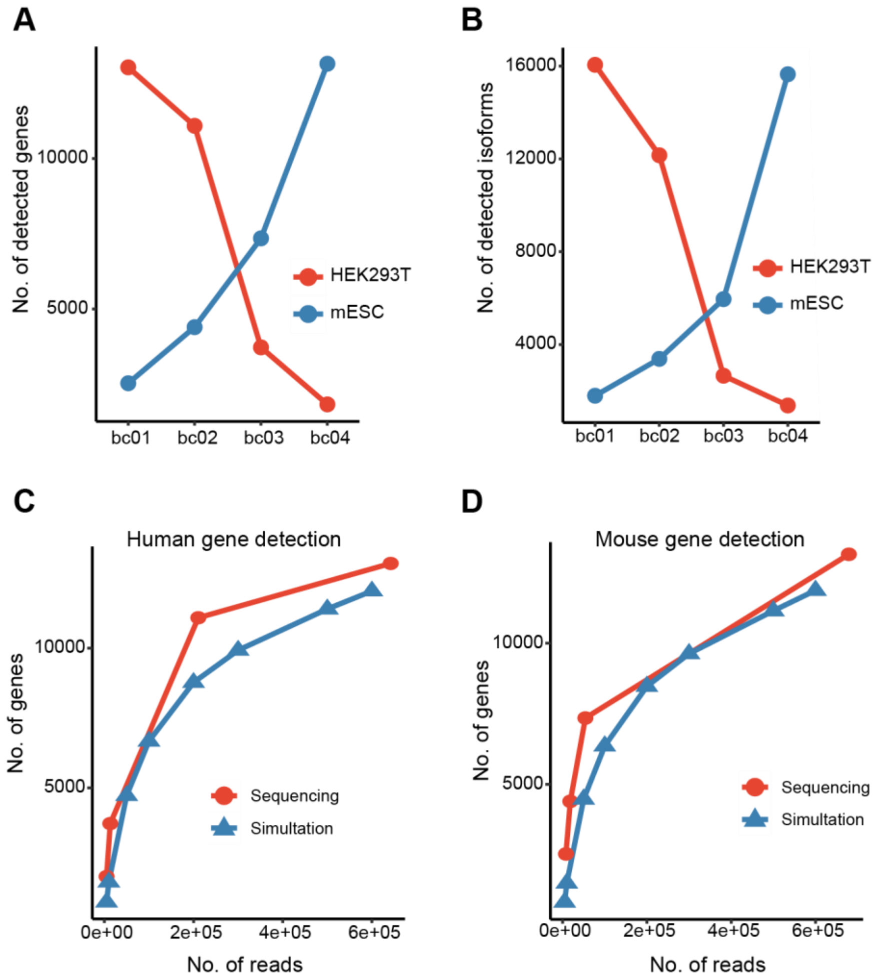


**Supplementary Figure 2. Gene and isoform detection between different total RNA inputs. (A)** Number of genes detected by RB-dRNA-seq in HEK293T and mESC cells using different total RNA input amounts. **(B)** Number of isoform detected by RB-dRNA-seq in HEK293T and mESC cells using different total RNA input amounts. **(C)** Gene detection sensitivity in HEK293T cells, comparing full RNA input amounts (x-axis) to downsampled read counts (simulated reduced sequencing depth). **(D)** Gene detection sensitivity in mESC cells, comparing full RNA input amounts (x-axis) to downsampled read counts (simulated reduced sequencing depth).

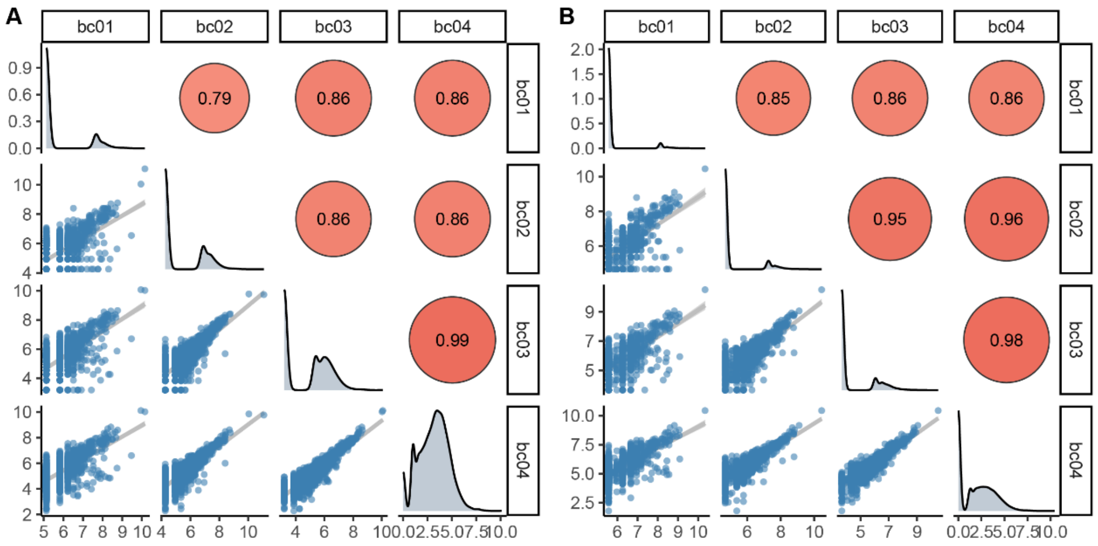


**Supplementary Figure 3. Gene expression and isoform expression correlation between different total RNA inputs.** Scatter plot showing gene expression **(A)** and isoform expression **(B)** correlation between different HEK293T total RNA inputs indicated by the RTA barcodes.

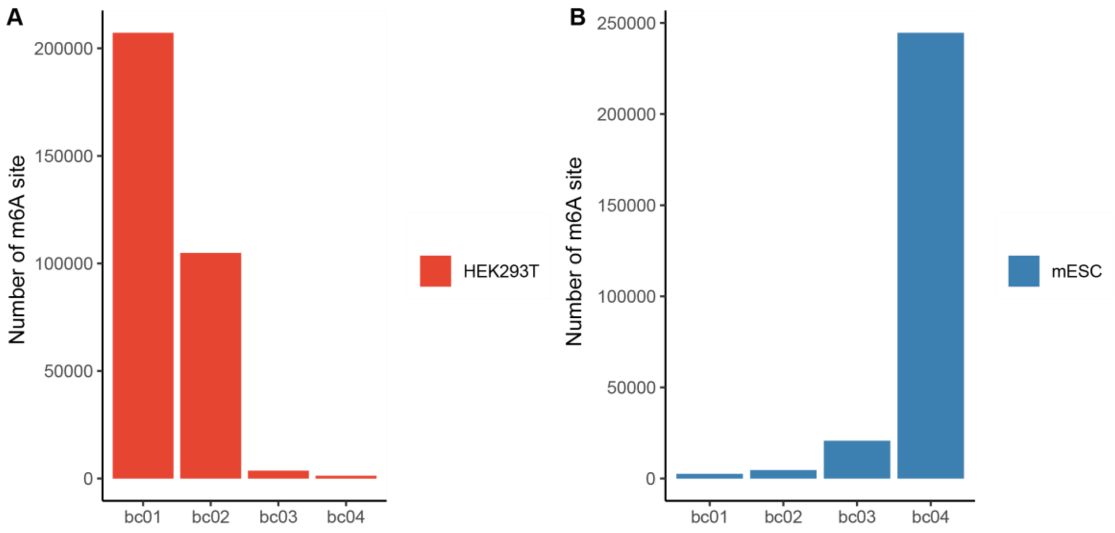


**Supplementary Figure 4. The m6A site detection between different total RNA inputs.** Number of m6A sites identified in HEK293T cells **(A)** and mESC cells **(B)** at different total RNA input amounts.

#### Supplementary Tables

| RTA Oligo | Sequence (5’->3’) |
|---|---|
| RTA-01_F | TCCAAGAAAGUUGUCGGUGUCUUUGUGTAGTAGGTTC |
| RTA-01_R | GAGGCGAGCGGTCAATTTTCACAAAGACACCGACAACTTTCTTGGATTTTTTTTTT |
| RTA-02_F | TCCUCGAUUCCGUUUGUAGUCGUCUGUTAGTAGGTTC |
| RTA-02_R | GAGGCGAGCGGTCAATTTTACAGACGACTACAAACGGAATCGAGGATTTTTTTTTT |
| RTA-03_F | TCCGAGUCUUGUGUCCCAGUUACCAGGTAGTAGGTTC |
| RTA-03_R | GAGGCGAGCGGTCAATTTTCCTGGTAACTGGGACACAAGACTCGGATTTTTTTTTT |
| RTA-04_F | TCCUUCGGAUUCUAUCGUGUUUCCCUATAGTAGGTTC |
| RTA-04_R | GAGGCGAGCGGTCAATTTTTAGGGAAACACGATAGAATCCGAAGGATTTTTTTTTT |
| RTA-05_F | TCCCUUGUCCAGGGUUUGUGUAACCUUTAGTAGGTTC |
| RTA-05_R | GAGGCGAGCGGTCAATTTTAAGGTTACACAAACCCTGGACAAGGGATTTTTTTTTT |
| RTA-06_F | TCCUUCUCGCAAAGGCAGAAAGUAGUCTAGTAGGTTC |
| RTA-06_R | GAGGCGAGCGGTCAATTTTGACTACTTTCTGCCTTTGCGAGAAGGATTTTTTTTTT |

**Supplementary Table 1. The sequences of RTA oligo and barcodes.** The underlined sequences in the table are RNA bases, and the un-underlined are DNA bases.

| | | RNA002 DNA 1BC | | RNA002 RNA 4BC | | RNA004 RNA 2BC | | | RNA004 RNA 4BC | | | RNA004 RNA 6BC | | |
|---|---|---|---|---|---|---|---|---|---|---|---|---|---|---|
| **Base Calling** | **Dorado Mode** | **fast** | **hac** | **fast** | **hac** | **fast** | **hac** | **sup** | **fast** | **hac** | **sup** | **fast** | **hac** | **sup** |
| | **Read Number** | 121117 | 121117 | 1058328 | 1058328 | 520748 | 520748 | 520748 | 227105 | 227105 | 227105 | 335407 | 335407 | 335407 |
| **Barcode Number** | **e-value<0.5** | 70961 | 77298 | 818769 | 814015 | 402096 | 467250 | 476341 | 180071 | 201441 | 202315 | 74020 | 221457 | 294094 |
| | **e-value<0.05** | 58371 | 66022 | 746711 | 745491 | 387516 | 457056 | 464532 | 172248 | 196345 | 196668 | 69217 | 213729 | 273000 |
| | **e-value<0.01** | 48982 | 59292 | 691930 | 699132 | 374754 | 447204 | 455285 | 165468 | 191408 | 192125 | 64975 | 207109 | 262940 |
| | **e-value<0.001** | 39325 | 50233 | 600402 | 629953 | 351448 | 427652 | 438738 | 153564 | 182632 | 184283 | 59311 | 196393 | 248176 |
| | **e-value<0.0001** | 28405 | 40014 | 513640 | 560898 | 325204 | 405015 | 419190 | 140226 | 172369 | 174823 | 52491 | 184648 | 232012 |
| **Barcode Number(%)** | **e-value<0.5** | 58.5888 | 63.8209 | 77.3644 | 76.9152 | 77.2151 | 89.7267 | 91.4725 | 79.2898 | 88.6995 | 89.0843 | 22.0687 | 66.0264 | 87.6827 |
| | **e-value<0.05** | 48.1939 | 54.5109 | 70.5557 | 70.4404 | 74.4153 | 87.7691 | 89.2048 | 75.8451 | 86.4556 | 86.5978 | 20.6367 | 63.7223 | 81.3937 |
| | **e-value<0.01** | 40.4419 | 48.9543 | 65.3795 | 66.0600 | 71.9646 | 85.8772 | 87.4290 | 72.8597 | 84.2817 | 84.5974 | 19.3720 | 61.7486 | 78.3943 |
| | **e-value<0.001** | 32.4686 | 41.4748 | 56.7312 | 59.5234 | 67.4891 | 82.1226 | 84.2515 | 67.6181 | 80.4174 | 81.1444 | 17.6833 | 58.5536 | 73.9925 |
| | **e-value<0.0001** | 23.4525 | 33.0375 | 48.5332 | 52.9985 | 62.4494 | 77.7756 | 80.4977 | 61.7450 | 75.8984 | 76.9789 | 15.6499 | 55.0519 | 69.1733 |
| **Barcode Accuracy(%)** | **e-value<0.5** | 97.3211 | 96.9960 | 98.2279 | 97.6693 | 99.0778 | 99.2208 | 99.1384 | 98.5561 | 98.8816 | 98.7861 | 96.2294 | 97.1850 | 94.1508 |
| | **e-value<0.05** | 99.3661 | 99.1215 | 99.5729 | 99.4814 | 99.7383 | 99.7650 | 99.7359 | 99.4531 | 99.4769 | 99.4442 | 98.7330 | 98.8074 | 98.8073 |
| | **e-value<0.01** | 99.8387 | 99.7841 | 99.8179 | 99.8090 | 99.8623 | 99.8656 | 99.8517 | 99.6561 | 99.6536 | 99.6627 | 98.9611 | 98.9720 | 99.0325 |
| | **e-value<0.001** | 99.9924 | 99.9622 | 99.8819 | 99.8878 | 99.8944 | 99.8885 | 99.8767 | 99.7805 | 99.7733 | 99.7791 | 99.0019 | 99.0580 | 99.1550 |
| | **e-value<0.0001** | 99.9894 | 99.9825 | 99.8923 | 99.9052 | 99.8961 | 99.8916 | 99.8791 | 99.8460 | 99.8248 | 99.8084 | 98.9903 | 99.0864 | 99.1871 |

**Supplementary Table 2. The results of barcode demultiplexing rate and demultiplexing accuracy.** Under different basecalling modes and e-value conditions. More detailed raw statistics for barcode demultiplexing can be found in the Supplementary Data appendix.

| | Barcode RTA | Total RNA input | Barcoded Reads | Detected Genes | Detected Isoform | Detected m6A site |
|---|---|---|---|---|---|---|
| **HEK203T** | Undemultiplexed | NA | 92510 | NA | NA | NA |
| | bc01 | 500ng | 642198 | 13034 | 16059 | 207214 |
| | bc02 | 100ng | 210667 | 11090 | 12159 | 104871 |
| | bc03 | 10ng | 13224 | 3724 | 2654 | 3670 |
| | bc04 | 1ng | 4900 | 1828 | 1371 | 1315 |
| **mESC** | Undemultiplexed | NA | 194808 | NA | NA | NA |
| | bc01 | 1ng | 8035 | 2531 | 1791 | 2563 |
| | bc02 | 10ng | 18286 | 4396 | 3381 | 4598 |
| | bc03 | 100ng | 54064 | 7349 | 5957 | 20762 |
| | bc04 | 500ng | 679180 | 13154 | 15652 | 244577 |

**Supplementary Table 3. The results of barcode demultiplexing and feature detection between different total RNA inputs.**

| | | Batch1 Read Count | Batch2 Read Count | Batch3 Read Count | Total Read Count |
|---|---|---|---|---|---|
| zygote | bc01 | 327 | 1616 | 515 | 8546 |
| zygote | bc02 | 631 | 2500 | 801 | |
| zygote | bc03 | 476 | 1341 | 339 | |
| 2-cell | bc04 | 2305 | 783 | 537 | 12252 |
| 2-cell | bc05 | 2966 | 581 | 434 | |
| 2-cell | bc06 | 3608 | 620 | 418 | |

**Supplementary Table 4. The results of barcode demultiplexing of low input embryos samples.**

| | **bc01** | **bc02** | **bc03** | **bc04** | **bc05** | **bc06** |
|---|---|---|---|---|---|---|
| **bc01** | 0 | 13 | 15 | 14 | 16 | 17 |
| **bc02** | 13 | 0 | 14 | 12 | 12 | 16 |
| **bc03** | 15 | 14 | 0 | 14 | 14 | 16 |
| **bc04** | 14 | 12 | 14 | 0 | 14 | 18 |
| **bc05** | 16 | 12 | 14 | 14 | 0 | 15 |
| **bc06** | 17 | 16 | 16 | 18 | 15 | 0 |

**Supplementary Table 5. The edit distance matrix between RNA barcodes.**
